# Division of labor between seed plant RAB GDI paralogs: insights from genetic analysis in *Arabidopsis thaliana*

**DOI:** 10.64898/2026.03.25.714218

**Authors:** Hana Soukupová, Fatima Cvrčková, Viktor Žárský, Michal Hála

## Abstract

**Background:** RAB Guanine Nucleotide Dissociation Inhibitors (RAB GDIs) are important vesicle transport regulators in eukaryotes, participating in the functional cycle of RAB GTPases by stabilizing their non-active GDP-conformation.

**Aims:** We address the importance of the three *Arabidopsis thaliana* RAB GDI paralogs by genetic and developmental analyses and put these results into the seed plants evolution context.

**Methods:** We use methods of genetics, microscopy and phylogenetics.

**Results:** Our genetic analyses of *Arabidopsis* T-DNA insertional mutants confirm recent CRISPR alleles data indicating lethality of double *gdi1 gdi2* mutants, and our microscopic data point to embryo development arrest in double mutant seeds. We also confirm the involvement of *GDI2* and *GDI3* in pollen tube growth. Moreover, our data show that *GDI1* also contributes to proper pollen function. Our phylogenetic analysis reveals independent diversification of RAB GDIs in Gymnosperms and Angiosperms, with early specialization of an Angiosperm reproduction-and gametophyte-related clade.

**Conclusions:** In *Arabidopsis*, RAB GDI1 and 2 are important for the vegetative growth while RAB GDI2 and 3 are vital for reproduction. Evolution of the RAB GDI family reflects the evolution of seed plants.

**Highlights:** RAB GDIs are vital for plant growth and reproduction and act redundantly. Even the low-transcribed RAB GDI1 isoform contributes to the proper pollen function. Two RAB GDI clades evolved in early Angiosperms.

## Introduction

RAB GDI is an important regulator of endomembrane vesicle trafficking, involved in the functional cycle of RAB GTPases, a branch of the larger superfamily of small GTPases. In plants, as in other organisms, RAB GTPases cycle between peripheral membrane attachment and cytoplasmic localization (reviewed e.g. in Ito and Uemura, 2022). RAB GTPases can be attached to membranes as peripheral membrane proteins due to the hydrophobic modification, double geranyl geranylation, occurring at the very C-terminus of the protein, where two cysteine motifs are usually localized, creating a prenylation site. Double geranyl geranylation of RAB GTPases is catalysed by the RAB geranylgeranyl transferase in concert with the RAB escort protein (REP; Hála et al., 2005; Hála et al., 2010).

RAB GTPases cycle between two conformational forms - a GDP-bound form, which is considered as inactive, and a GTP-bound form, which is considered to be the active form – acting as a molecular switch regulating downstream interacting protein effectors. Binding of GTP changes the conformation of the Switch II region and enables interaction with effector proteins. RAB GTPases are very slow hydrolases that need the GTPase activating protein (GAP) to speed up the conversion from the GTP-bound state to the GDP-bound state. Conversely, the RAB GTPases can be recharged by the action of the Guanosine nucleotide exchange factor (GEF; Ito and Uemura, 2022). Following GTP hydrolysis, the inactive GDP-bound RAB GTPase is extracted from the membrane by RAB GDI and transported back to the cytoplasm for the next round of activation. Since the interaction between RAB GTPase and RAB GDI is relatively strong, the dissociation of the complex is facilitated by the action of the GDI displacement factor (GDF; Bahk et al., 2009).

RAB GDIs in plants were first characterized by Zarsky et al. (1997), Andreeva et al (1997) and Ueda et al. (1996,1998) in experiments complementing loss of the unique yeast RAB GDI Sec19p by *Arabidopsis* homologs. This experiment also demonstrated functional conservation of RAB GDIs among kingdoms. RAB GDIs are closely related to REPs (Hála et al., 2005) as both families share the typical structure comprising two domains. The Domain I recognizes the RAB GTPase itself, and the Domain II provides a binding platform for the prenylation tail (Pylypenko et al., 2006). Both proteins bind RAB GTPases promiscuously. In the *Arabidopsis* genome, there are 57 RAB GTPases but only three GDIs and only one REP (Rutherford and Moore, 2002; Hála et al., 2005).

RAB GDIs may be considered as house-keeping proteins. However, they have been documented to exist in multiple isoforms with tissue- or cell type-specific expression in metazoans (e.g. Yang et al., 1994; Benhar et al., 1997). In plants, RAB GDI isoforms have been implicated in the responses to various stress conditions. Using GWAS analysis, Amangeldyieva et al. (2025), identified soybean RAB GDI as one of two downregulated genes in accessions with increased resistance to drought stress. Similarly, wild tomato RAB GDI, SchRABGDI1, was shown to be involved in the response to the salinity stress and its heterologous expression in *Arabidopsis* increased salt resistance (Martín-Davison et al., 2017). Tobacco RAB GDI was also shown to be one of the genes conferring enhanced tolerance to aluminium ions and overexpression of *NtGDI1* in yeast increased their tolerance to aluminium in the medium (Ezaki et al., 2005). Finally, maize homolog of RAB GDI-alpha was shown to be crucial for the penetration of virus to the plant cells. A recessive *RAB GDI* allele caused by helitron transposon insertion resulting in protein sequence changes due to alternative splicing confers resistance to the virus, which cannot bind the mutant protein (Liu et al., 2020).

Importance of RAB GDI proteins for plant ontogeny was addressed several times. In a systematic screen of CRISPR/Cas9-generated mutants in various subsets of the 6 predicted rice RAB GDI paralogs, Shad et al (2023) observed that plants lacking GDI1 exhibit significantly shorter shoots and shorter grains, while inactivation of some other RAB GDIs led to lethality and no multiple mutants were detected. Recently, the role of RAB GDIs in embryogenesis of *Arabidopsis thaliana* was studied using CRISPR/Cas9 based mutants. It was shown that RAB *GDI1* and *GDI2* are crucial for embryogenesis, while the double mutation is embryo-lethal (Yin et al., 2026). In another study, the importance of the RAB *GDI2* and *GDI3* for the pollen development and the pollen tube growth was documented by showing that there is no transmission of double *gdi2/gdi3* mutation through the pollen (Wu et al., 2025).

In this report, we investigate an until-now-uncharacterized set of T-DNA loss of function mutant alleles of all three *Arabidopsis* RAB GDI paralogs. Besides confirming the reported observations from CRISPR mutants (Wu et al., 2025), we show that RAB GDI1 also contributes to the pollen function. Our phylogenetic analysis indicates early diversification of two Angiosperm RAB GDI clades that adopted specialized functions as inferred from their differential expression in various tissues. In addition, an unrelated duplication event was identified in the Pinales, indicating possible parallel diversification in the Gymnosperms.

## Methods

### Plant materials and growth conditions

Seeds of *Arabidopsis thaliana* were surface sterilised, imbibed in sterile water for 2 days at 4°C and germinated on ½ MS media supplemented with 1% sucrose, vitamin mixture, and 0,8% plant agar (Duchefa Biochemie) 22 °C, 16/8 light/dark regime. In this study we used loss of function (LOF) T-DNA insertional mutants in *GDI1* - At2g44100 (*gdi1-1*, obtained from Klaus Palme, MPI Cologne, as L59, with T-DNA insertion very close to that in the GABIseq_180B061 line), GDI2 - At3g59920 (*gdi2-1*, SALK_064619) and GDI3 - At5g09550 (*gdi3-1*, FLAG_109C08,109E12). Localization of the insertions is described in Fig.1. For simplicity, the indicated mutant alleles will be further referred to as *gdi1, gdi2* and *gdi3*.

**Fig. 1.**
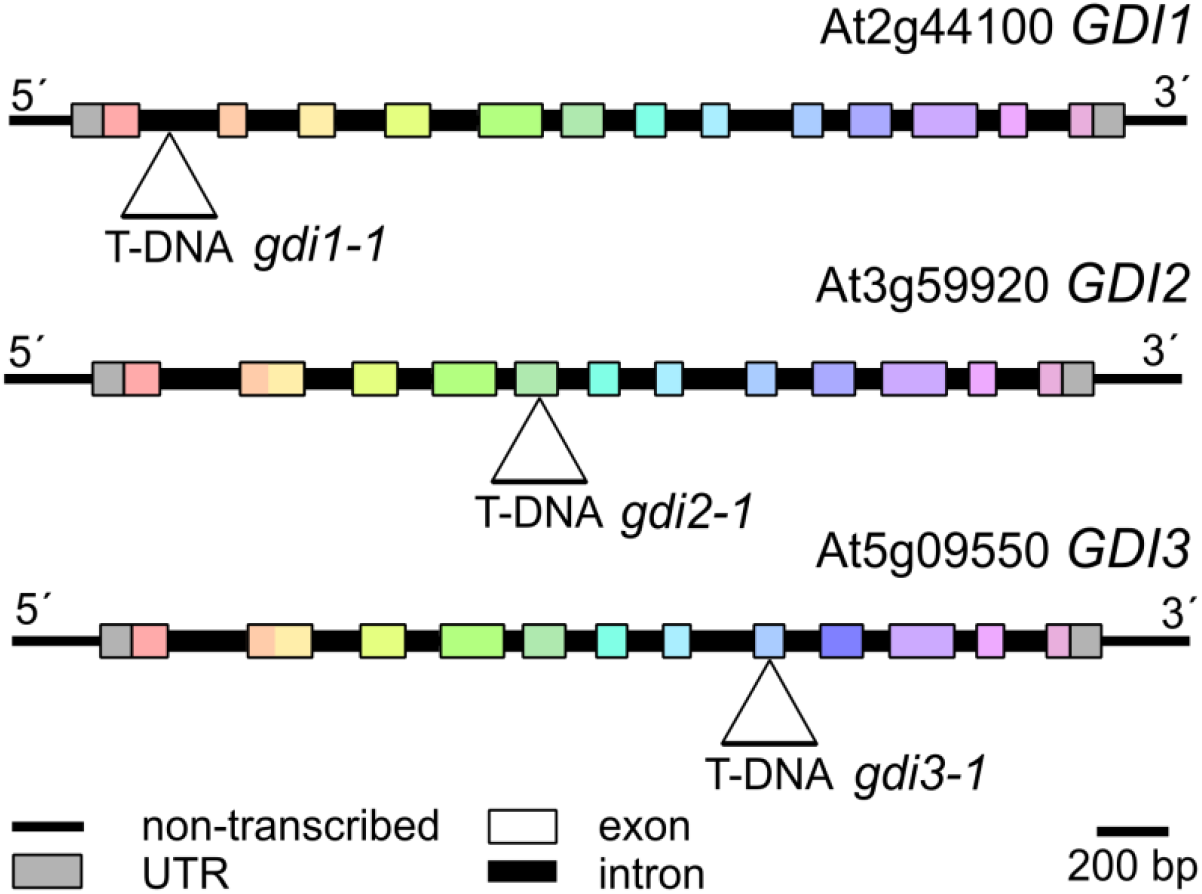
Map of the three *A. thaliana* GDI loci with positions of T-DNA insertions indicated. Homologous exons are shown in the same color.

Selection plates containing sulfadiazine (100 mg/l) were used for selection of *gdi1*, kanamycin (50 mg/l) for *gdi2*, and PPT/Basta (20 mg/l) for *gdi3* mutant plants. Young seedlings were transferred to pellets (Jiffy Products International) and moved to *ex vitro* growth condition. The genomic DNA was extracted from young seedlings using DNAzol (Invitrogen) reagent according to the manufacturer. PCR analyses were performed using primer combinations listed in the Supplementary table 1.

### Transmission ratio and seed set analyses

For transmission ratio determination, F1 progeny of crossed plants was evaluated by genotyping all seeds obtained from manually pollinated flowers (even in case of identical mother and father plant genotypes, i.e., progeny from selfing was not analysed in these experiments). Each crossing was performed on at least three mother plants.

Significance of segregation ratios deviation from Mendelian ones was determined using the goodness of fit Chi square test using an online calculator (https://www.socscistatistics.com/).

For seed set efficiency determination, siliques were divided at half of their length and seeds in the apical (stigma) and basal part were counted separately, while also recording the total seed count.

### Microscopic observations

Developing seeds were dissected from young siliques using a fine needle under the stereo microscope Leica S6D. Pictures of siliques with visible phenotypic variation of seeds (green vs. white) were taken with the attached camera Canon PowerShot S70. Seeds from several developmental stages of young siliques were cleared by fresh 0,5M KOH and embryos in transparent seeds were directly observed on the Olympus BX51 microscope equipped with the camera Olympus DP74.

### Phylogenetic tree construction

Angiosperm and *Ginkgo biloba* GDI-encoding genes were identified in annotated genomes and species-specific GenBank Refseq sections by a combination of keyword-based searches and exhaustive BLAST searches with the *A. thaliana* GDI1 protein sequence query. For several other Gymnosperm species, similar BLAST searches were performed in species-specific transcriptome shotgun assemblies (TSA), and for *Picea sitchensis*, two previously reported GDI sequences (Wu et al., 2025) have been included. The full list of genes included in the phylogenetic tree is provided in Supplementary table 2. Since preliminary attempts to construct a phylogenetic tree based on amino acid sequences yielded poor statistical support due to the near-perfect conservation of large portions of the GDI proteins, nucleotide sequences of the GDI-encoding ORFs were retrieved for all genes and aligned manually in BioEdit (Hall, 1999) version 7.7.1 employing the “toggle translation” mode (i.e. using the well conserved protein sequences as a seed for nucleotide alignment). The resulting nucleotide alignment has been used to construct a maximum likelihood phylogenetic tree in MEGA 12 (Kumar et al., 2024), using standard bootstrap validation with default settings and 500 bootstrap cycles, and excluding all gap-containing columns.

## Results

### Genetic analysis of the three Arabidopsis RAB GDI isoforms

The efficiency of LOF mutation transmission in backcrosses is a good indicator of the importance of the gene for the male or female gametophyte. We used various combinations of the three *RAB GDI* alleles as either pollen acceptors or donors (i.e. “mother” or “father”) and evaluated the transmission rate of the mutations. Results are summarized in Table 1.

**Table 1.**
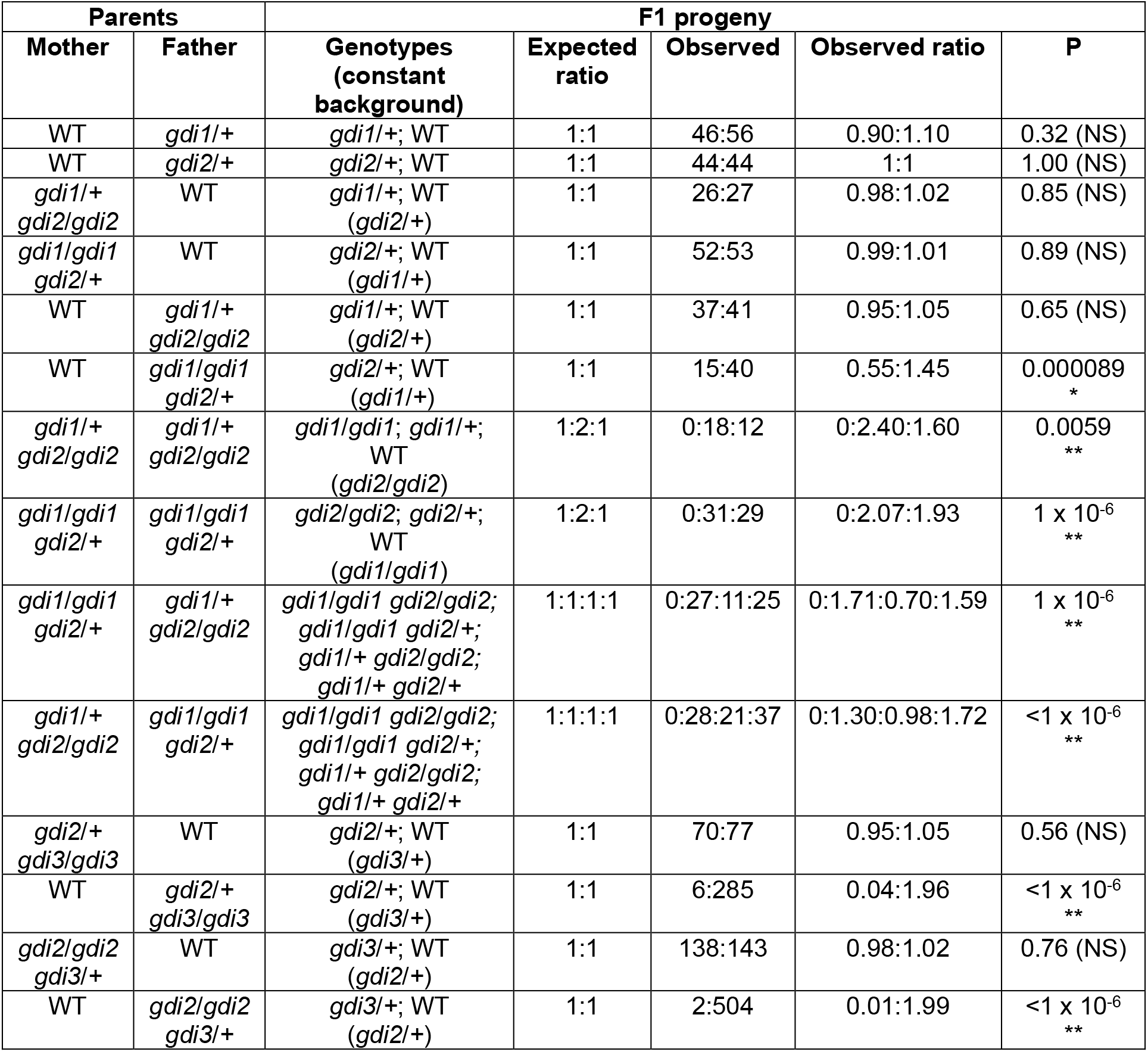
Genetic analysis of *Arabidopsis* RAB GDI mutants. Numbers of plants with expected genotypes identified in the F1 progeny are given in the Observed column. P-values were calculated by goodness of fit chi square test (https://www.socscistatistics.com/).

| Parents |  | F1 progeny |  |  |  |  |
| --- | --- | --- | --- | --- | --- | --- |
| Mother | Father | Genotypes (constant background) | Expected ratio | Observed | Observed ratio | P |
| WT | *gdi1/+* | *gdi1/+*; WT | 1:1 | 46:56 | 0.90:1.10 | 0.32 (NS) |
| WT | *gdi2/+* | *gdi2/+*; WT | 1:1 | 44:44 | 1:1 | 1.00 (NS) |
| *gdi1/+*<br>*gdi2/gdi2* | WT | *gdi1/+*; WT<br>(*gdi2/+*) | 1:1 | 26:27 | 0.98:1.02 | 0.85 (NS) |
| *gdi1/gdi1*<br>*gdi2/+* | WT | *gdi2/+*; WT<br>(*gdi1/+*) | 1:1 | 52:53 | 0.99:1.01 | 0.89 (NS) |
| WT | *gdi1/+*<br>*gdi2/gdi2* | *gdi1/+*; WT<br>(*gdi2/+*) | 1:1 | 37:41 | 0.95:1.05 | 0.65 (NS) |
| WT | *gdi1/gdi1*<br>*gdi2/+* | *gdi2/+*; WT<br>(*gdi1/+*) | 1:1 | 15:40 | 0.55:1.45 | 0.000089<br>* |
| *gdi1/+*<br>*gdi2/gdi2* | *gdi1/+*<br>*gdi2/gdi2* | *gdi1/gdi1*; *gdi1/+*;<br>WT<br>(*gdi2/gdi2*) | 1:2:1 | 0:18:12 | 0:2.40:1.60 | 0.0059<br>** |
| *gdi1/gdi1*<br>*gdi2/+* | *gdi1/gdi1*<br>*gdi2/+* | *gdi2/gdi2*; *gdi2/+*;<br>WT<br>(*gdi1/gdi1*) | 1:2:1 | 0:31:29 | 0:2.07:1.93 | 1 x 10<sup>-6</sup><br>** |
| *gdi1/gdi1*<br>*gdi2/+* | *gdi1/+*<br>*gdi2/gdi2* | *gdi1/gdi1* *gdi2/gdi2*;<br>*gdi1/gdi1* *gdi2/+*;<br>*gdi1/+* *gdi2/gdi2*;<br>*gdi1/+* *gdi2/+* | 1:1:1:1 | 0:27:11:25 | 0:1.71:0.70:1.59 | 1 x 10<sup>-6</sup><br>** |
| *gdi1/+*<br>*gdi2/gdi2* | *gdi1/gdi1*<br>*gdi2/+* | *gdi1/gdi1* *gdi2/gdi2*;<br>*gdi1/gdi1* *gdi2/+*;<br>*gdi1/+* *gdi2/gdi2*;<br>*gdi1/+* *gdi2/+* | 1:1:1:1 | 0:28:21:37 | 0:1.30:0.98:1.72 | <1 x 10<sup>-6</sup><br>** |
| *gdi2/+*<br>*gdi3/gdi3* | WT | *gdi2/+*; WT<br>(*gdi3/+*) | 1:1 | 70:77 | 0.95:1.05 | 0.56 (NS) |
| WT | *gdi2/+*<br>*gdi3/gdi3* | *gdi2/+*; WT<br>(*gdi3/+*) | 1:1 | 6:285 | 0.04:1.96 | <1 x 10<sup>-6</sup><br>** |
| *gdi2/gdi2*<br>*gdi3/+* | WT | *gdi3/+*; WT<br>(*gdi2/+*) | 1:1 | 138:143 | 0.98:1.02 | 0.76 (NS) |
| WT | *gdi2/gdi2*<br>*gdi3/+* | *gdi3/+*; WT<br>(*gdi2/+*) | 1:1 | 2:504 | 0.01:1.99 | <1 x 10<sup>-6</sup><br>** |

We first focused on the *GDI1* and *GDI2* genes that are expressed in a wide variety of vegetative tissues according to publicly available transcriptome data, while GDI3 transcript is only detected in pollen and possibly anthers (Waese et al., 2017). Single homozygous LOF mutations of either *GDI1* or *GDI2* had no effect upon transmission during backcrossing as the observed transmission rate was not significantly different from the expected 1:1 (WT: heterozygotes) ratio. We then backcrossed combined double mutants in the hemizygous state (i.e. one of the mutations was homozygous and the other heterozygous). While the *gdi2* allele was transmitted as efficiently as WT from mother plants harbouring a homozygous *gdi1* mutation, significantly fewer F1 plants carrying *gdi2* were recovered in the opposite backcross direction, with a transmission ratio of 0.55:1.45 instead of 1:1, indicating decreased pollination efficiency of double mutant *gdi1 gdi2* pollen (which might still contain some paternal *GDI2* gene product transmitted through meiosis) in direct competition with *gdi1GDI2* pollen while no such effect was observed in *gdi2* vs. *GDI2* pollen competition (see above). However, the opposite combination, i.e. transmission of *gdi1* from a heterozygote in a homozygous *gdi2* background, yielded transmission ratios not significantly differing from the theoretical 1:1 ratio regardless of the backcross direction. This observation suggests that *GDI1* and *GDI2* contribute to the male gametophyte function unequally, with *GDI2* being more critically required for efficient fertilization. This may be related to their different expression patterns - while *GDI2* exhibits high transcript levels in both vegetative tissues and pollen, *GDI1* expression in pollen is negligible (Waese et al., 2017). Attempts to obtain double homozygous *gdi1/gdi1 gdi2/gdi2* mutants by crossing all possible hemizygote combinations (i.e. *gdi1* heterozygotes in the *gdi2* homozygous background and vice versa) in both directions failed, and the resulting F1 progeny often harboured fewer heterozygotes than expected, again indicating that *GDI1* and *GDI2* share a partially redundant essential function in fertilization or early development.

We then performed analogous backcross analyses involving the LOF allele of *GDI3*. We readily obtained heterozygous *gdi3* plants harbouring homozygous *gdi2* and vice versa (i.e. heterozygous *GDI2* in a homozygous *gdi3* background). In both cases, when the mutant plants were acceptors of WT pollen, the transmission ratios were close to expected 1:1, but simultaneous transmission of *gdi2* and *gdi3* mutations through pollen was dramatically reduced from the expected 1:1 ratio to 0.04:1.96 (*gdi2/+gdi3/+*: *GDI2/+gdi3/+* in F1 generation) or 0.01:1.99 (*gdi2/+gdi3/+*: *gdi2/+GDI3/*+ in F1 generation), respectively (Table 1). Upon selfing however, we were able to obtain double homozygous mutants with a very low frequency. This points to the importance of both pollen-expressed GDI paralogs for male gametophyte function.

### RAB GDI double mutants are embryonic lethal

The absence of double homozygous *gdi1 gdi2* mutants raises the question whether such mutants are produced at all and if so, at what developmental stage is their development arrested. To answer this, we analysed siliques of selfed *gdi1* heterozygotes in a *gdi2* homozygous background. Normal segregation ratio predicts one quarter of double homozygotes, which were however absent from our genetic analyses (see above). Upon selfing the hemizygous plants, approximately one quarter of positions in the silique is occupied by aberrant seeds exhibiting pale or milky colour and smaller size (Fig. 2A).

**Fig. 2.**
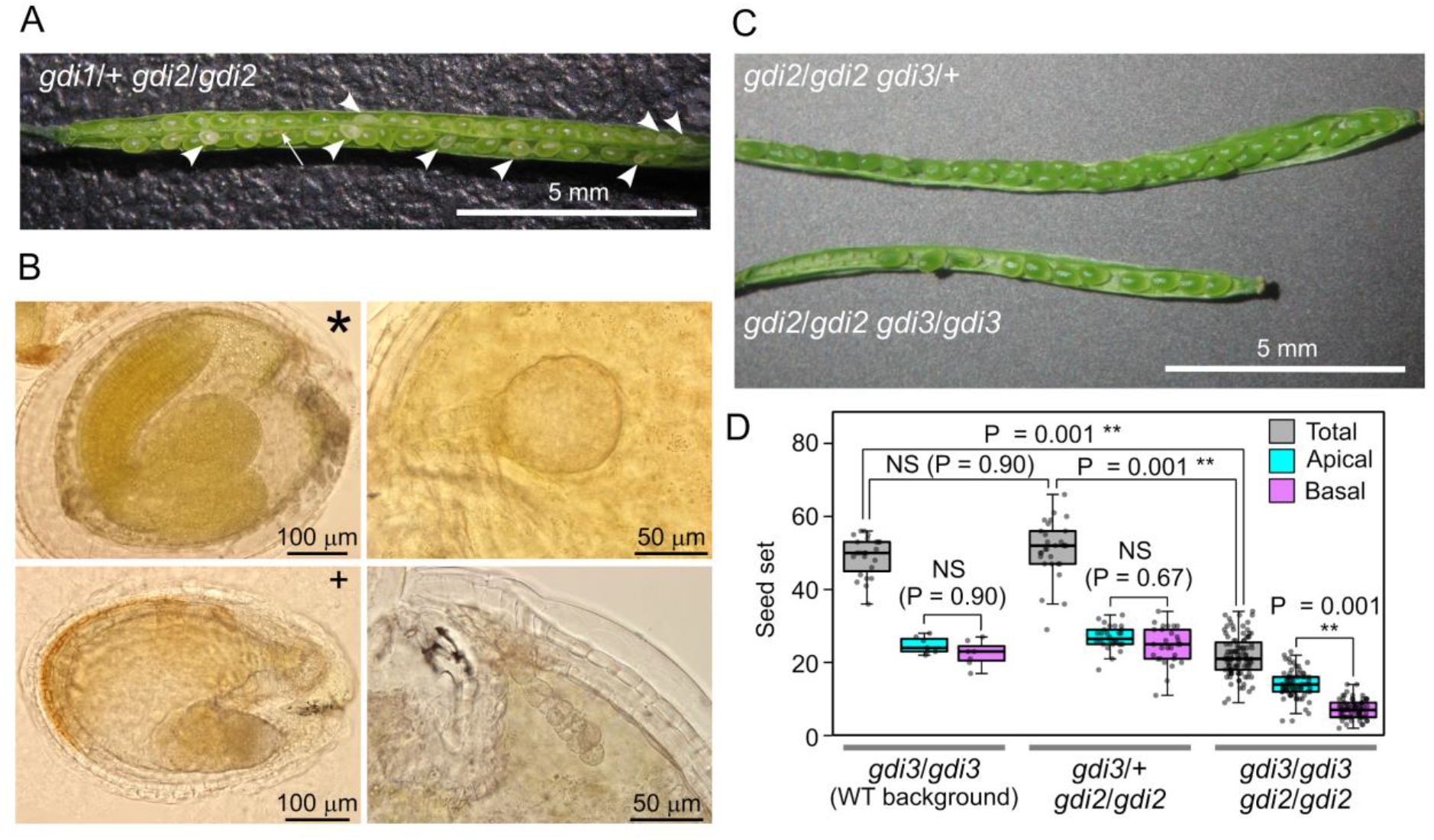
Phenotypic deviations of *Arabidopsis* RAB GDI double mutants. (A) A typical silique of a plant carrying a heterozygous *gdi1* mutation in a *gdi2* mutant background. Arrowheads denote defective (pale/milky) seeds; the arrow indicates a seed aborted at an early developmental stage; stigma is oriented to the right. (B) Examples of defective seeds from the same genotype, with developmentally arrested or defective embryos, shown alongside a healthy embryo (marked by an asterisk). All embryos are of similar age except the defective one marked by a cross, which is from a somewhat more advanced silique. (C) Typical siliques of plants harbouring the *gdi3* mutation in either heterozygous or homozygous state in a *gdi2* mutant background; stigma is oriented to the right. (D) Comparison of seed set in homo- or heterozygous *gdi3* mutants in wild type or *gdi2* mutant background, evaluated for complete siliques (“total”) and their apical (stigma-oriented) and basal halves, respectively. Statistical significance (one-way ANOVA, Tukey HSD) is shown for selected combinations. At least 24 siliques were evaluated per genotype.

Although we could not verify the genotype of these collapsing seed initials by PCR analysis, we hypothesize that these seeds contain double mutant embryos. Further microscopic characterization revealed that the aberrant seeds contain embryos arrested at the globular developmental stage or callus-like embryonic tissues, and occasionally larger malformed embryos (Fig. 2B), supporting the hypothesis that double homozygous *gdi1 gdi2* mutants die during early postzygotic development.

The situation was different in plants harbouring a combination of the *gdi2* and *gdi3* mutations. Although simultaneous transmission of both mutant alleles through pollen was very rare (see above), the double homozygous plants are viable. Upon selfing however, they form siliques that are significantly shorter than those of hemizygous *(gdi2/gdi2 gdi3/+)* plants (Fig.2C). The double mutant siliques contain significantly fewer seeds than those of either WT or hemizygous plants. Moreover, the efficiency of ovule fertilization in these double mutants is significantly lower in the basal half of the silique compared to the part adjacent to the stigma, indicating that mutant pollen tubes have an obvious difficulty in reaching the bottom part of the carpel, i.e. an obvious male gametophytic defect (Fig.2D).

### Two GDI clades are an Angiosperm apomorphy

Typically, plant genomes encode multiple GDI isoforms and this observation has been recently interpreted as indicating an ancient gene duplication (Wu et al 2025), although detailed phylogenetic analysis was not yet conducted. To clarify phylogenetic relationships within the seed plant GDI gene family, we performed exhaustive sequence similarity searches of representative genomes. Form Angiosperms, the selected eudicot species included *A. thaliana*, poplar (*Populus trichocarpa*), barrel medic (*Medicago truncatula*), soybean (*Glycine max*), and grapevine (*Vitis vinifera*), while monocots were represented by three grasses - maize (*Zea mays*), rice (*Oryza sativa*) and sorghum (*Sorghum bicolor*). We also included *Amborella trichopoda*, representing a basal Angiosperm clade considered sister to both monocots and dicots (Shi and Van de Peer, 2026). Gymnosperms were represented by *Gingko biloba*, yew (*Taxus sinensis*), and four Pinaceae representatives - a pine (*Pinus taeda*), two spruce species (*Picea abies* and *P. sitchensis*), and a larch (*Larix siberica*).

All the examined genomes encoded between two and five RAB GDI paralogs, with the exception of *G. biloba*, where only one RAB GDI-encoding gene was found (Supplementary Table 2). As the protein sequences of RAB GDIs are very similar, we performed the phylogenetic analysis on coding DNA sequences, which provided stronger phylogenetic signals, allowing for construction of a robust phylogenetic tree (Figure 3). The analysed RAB GDI sequences clearly segregated into three clades, one of containing all Gymnosperm representatives, while each of the remaining two clades, further referred to as Angiosperm clade 1 and Angiosperm clade 2, contained at least one representative from each of the examined Angiosperm species. With a single exception (concerning the position of one *Amborella* sequence), the three main clades had very strong bootstrap support. This suggests that the two Angiosperm clades originated by a gene duplication already in the common Angiosperm ancestor.

**Fig. 3.**
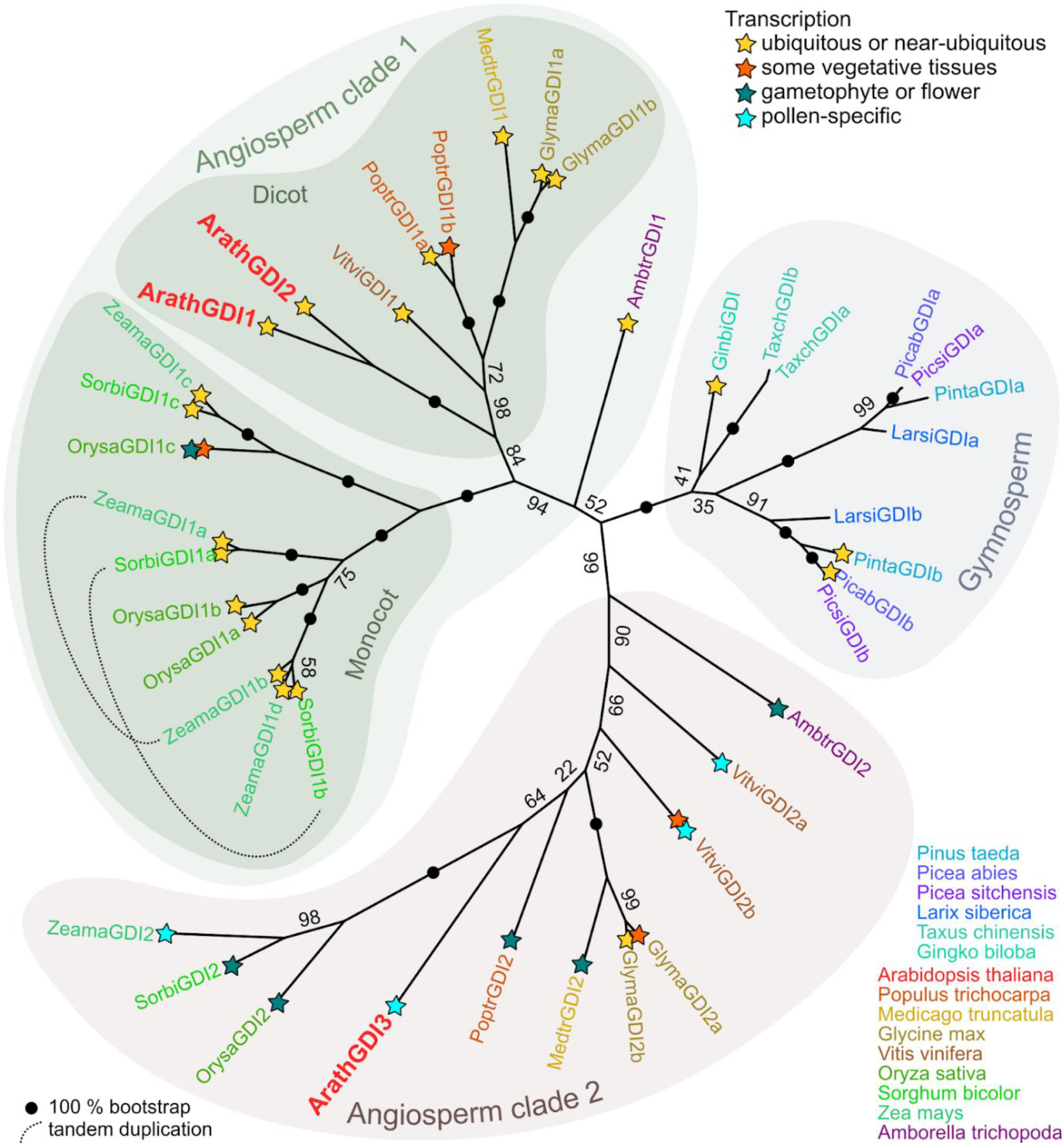
Phylogeny of seed plants RAB GDIs. A maximum likelihood phylogenetic tree reconstructed from an alignment of coding nucleotide sequences as described in Methods is shown, and symbols denoting the overall transcription pattern are provided for genes with available transcriptome data. For a full list, terminology and description of the included loci, and the transcriptome evidence see Supplementary table 2.

Further duplications took place subsequently in all three clades. Within the Angiosperm clade 1, two deep-branching families of paralogs are present in all studied monocots, and in one of them, subsequent gene duplications took place. Interestingly, in both maize and *Sorghum*, these include a tandem duplication that may have predated the divergence of lineages leading to these two grasses. Within Clade 2, there are generally fewer paralogs, and gene duplications were documented only in soybean and grapevine, with the former apparently recent but the latter probably ancient. In the Gymnosperms, all studied species except *Ginkgo* had two Rab GDI paralogs that could be attributed to a gene duplication event that took place before diversification of Pinaceae and an independent recent duplication in *Taxus*.

Some insights into the functional diversification of GDI paralogs could be gained from publicly available transcriptome data (Figure 3, Supplementary Table 2). In general, members of the Angiosperm clade 1 are typically expressed ubiquitously or in a wide variety of vegetative tissues. Eventually, there are genes expressed predominantly in specific tissues. It concerns, for example, the clade containing OrysaGDI1c, which is reported to be predominantly expressed in the aleurone (Rice expression database, https://ngdc.cncb.ac.cn/red/). On the other hand, the Clade 2 genes tend to be expressed predominantly, and in some cases exclusively, in reproductive organs or in the gametophyte. In several cases, including ArathGDI3, a clade 2 isoform appears to be pollen-specific, and in at least one case (*M. truncatula*) the observed expression pattern does not exclude pollen specificity since only male flower or anther, but not pollen, RNA data are available. The available transcriptome data do not indicate a similarly consistent pattern among Gymnosperm GDIs, although it is conspicuous that in at least two Pinaceae species one of the two early-diverging clades is represented by multiple ESTs and detected in a variety of tissues, while the other has only a few ESTs and remains below detectable levels in vegetative tissue transcriptomes.

### Discussion

Rab GDIs are important housekeeping proteins contributing via RAB GTPases to the overall dynamicity of the endomembrane trafficking in eukaryotes. Generally, their function is connected with extraction of inactive GDP-bound RAB GTPases from the membranes, though “moonlighting” roles were reported in some cases (McLauchlan et al., 1998). In this report we address several general aspects concerning RAB GDI functionality in *Arabidopsis* using primarily genetic analysis of three *Arabidopsis* GDIs LOF mutants.

Rab GDIs belong to a very conserved eukaryotic protein family. Sometimes they are grouped with REPs to the REP-GDI superfamily on the basis of sequence and structure similarity, although the two types of proteins have different functions (Hála et al, 2005). The RAB GDI sequences are highly conserved across seed plant species, typically exceeding 75% protein sequence identity. That is why we preferred to use nucleotide sequences to construct our phylogenetic tree. Despite the RAB GDI sequence conservation, there are alternative, usually longer, splice variants reported (e.g. the maize Zm00001eb349250_T001 sequence at http://phytozome-next.jgi.doe.gov). This suggests that, however conserved the sequences are, there is a possibility to adopt new sequence features for possible new functions. This is also exemplified by the alternative splicing of maize RAB GDI exon 10 upon insertion of the helitron transpozone, leading to a RAB GDI splicing variant, which is not recruited by viral protein and thus causes dominant negative resistance to maize rough dwarf disease (Liu et al., 2020).

Most of the seed plants’ genomes encode more than one RAB GDI isoform, raising questions about redundancy and sub-functionalization of these paralogs, which might be first approached using tissue or developmental stage-specific public transcriptomic data. As pollen tubes are among the most secretory active cell types, it is interesting to look there. The data show that most abundant RAB GDI isoforms on the mRNA level in *Arabidopsis* pollen are GDI2 and GDI3, and recently published functional data obtained using CRISPR/Cas9 mutants confirm pollen roles of these two paralogs (Wu et al, 2025). Our data obtained on not yet studied T-DNA insertion alleles are in accord with these findings, showing tremendously decreased transmission of *gdi2 gdi3* double mutation through the pollen. The remaining isoform, GDI1, exhibits barely detectable expression levels in pollen. Proteomic analysis of *Arabidopsis* pollen identified only one peptide that can be assigned to GDI1 (Grobei et al., 2009). Similarly, analysis of GDI1 transcript level in pollen revealed only a very weak signal (Yin et al., 2025). Taken together, GDI1 does not seem to be the important isoform for the pollen function from this quantitative point of view. Yet, in contrast to conclusions of Wu et al. (2025) our data clearly show that also GDI1 has a role in pollen function, as its loss in hemizygous double mutants (*gdi1*/*gdi1 gdi2*/+) significantly decreases the fertilization efficiency of *gdi2* pollen tubes compared to pollen produced by single homozygous *gdi2* mutant plants. As GDIs are largely promiscuous in respect to interactions with RAB GTPases and no specificity has been reported till now, it is possible that loss of GDI2 may be compensated by the GDI1 paralog whose expression might be upregulated in *gdi2* pollen. Outside plants, some functional specialization was reported between alpha and beta GDI isoforms in mice brain (Benhar et al., 1997). The importance of GDI1 and RAB GDIs in pollen generally might also be connected with not yet discovered interaction outside the RAB GTPase family. In mammals for example, RAB GDI was reported to interact with a membrane-bound HSP90 complex (Chen and Balch, 2006). It is also possible that the specificity of the interaction may be brought by the GDFs recognizing different complexes of RAB GTPases with GDIs (Dirac-Svejstrup et al., 1997).

Our phylogenetic analysis clearly shows two specific separate deep GDI branches in Angiosperms that appear to be conserved from the basal *Amborella* through dicots and at least a sizeable portion of monocot lineages (though we only analysed a set of grass genomes). An independent diversification of two RAB GDI clades took place also in the Gymnosperms at or below the base of Pinales. Unfortunately, the position of the two gymnosperm plants outside Pinales that were included in our analysis, namely *Ginkgo* and *Taxus*, with respect to these two branches could not be resolved because the well conserved gene sequences did not carry sufficient phylogenetic signals. Transcriptomic data from *Arabidopsis*, rice and maize suggest that one Angiosperm branch – including RAB GDI3 of *Arabidopsis* - contains isoforms highly expressed in reproductive organs, gametophytes, and in some cases exclusively in pollen. Pollen tube is a specific organ that demands high overall RAB GTPases activity as a component of membrane trafficking machinery sustaining rapid pollen tube growth. Interestingly, the Gymnosperms also show similar independent diversification in the common ancestor of Pinales. Conifers independently evolved pollen tubes as a tool to deliver immotile sperm cells to ovules for fertilization (siphonogamy; Lora et al., 2016). These pollen tubes are slowly growing, often branched and do not overcome long distances. It may be thus tempting to speculate that the second GDI clade in Gymnosperms might be specialized for siphonogamy, since one of the clades is dramatically underrepresented in sporophytic transcriptomes, only one RAB GDI is found in in non-siphonogamous *Gingko*. A very recent RAB GDI gene duplication took place also independently in siphonogamous Chinese yew. Unfortunately, there are no public transcriptome data from Gymnosperm pollen and thus no experimental evidence for this hypothesis, which would deserve to be tested by GDI-specific RT-PCR in some conifer and its pollen. However, we can safely conclude that in both Angiosperms and Gymnosperms, the available gene expression pattern data suggest specialization of individual Rab GDI clades, although in the Gymnosperm case we can only speculate about the nature of such specialization.

## Abbreviations

GAP: GTPase Activating Protein
GEF: Guanosine Nucleotide Exchange Factor
GDF: GDI Displacement Factor
GDI: Guanine Nucleotide Dissociation Inhibitors
LOF: Loss of Function
REP: RAB Escort Protein
TSA: Transcriptome Shotgun Assemblies

## Supplementary files

**Supplementary table 1. Primers for PCR genotyping analyses of RAB GDI alleles**

**Supplementary table 2. List of RAB GDI accessions used for the phylogenetic tree construction**.

## Acknowledgements

This work was supported by the project TowArds Next GENeration Crops, reg. no. CZ.02.01.01/00/22_008/0004581 of the ERDF Programme Johannes Amos Comenius. Authors also thank Dr. Klaus Palme and Dr. Frieder Bischoff for the mutant line L59, which later became GABIseq_180B061.

## Data and materials availability statement

All the data used for the paper preparation are presented in the main text body or in supplements. All the T DNA mutant lines used in the paper come from public collections and all the sequences were retrieved from open databases as mentioned in the text or supplements.

## Declaration of ethical and legal standards compliance

This work was prepared in accordance with the rules concerning the work with GMO plants.

