## Supplementary table 1 for "Division of labor between seed plant RAB GDI paralogs: insights from genetic analysis in *Arabidopsis thaliana*"

**Supplementary Table 1. Primers for PCR genotyping analyses of RAB GDI alleles**

| | |
|---|---|
| *GDI1-1* | |
| GDI1-Fw | GGAGATTTTCAATACTCTTAATTGC |
| GDI1-Rv | GAAGTACCTAATTGGAATATAGATG |
| RB-TDNA-GABIKat | AGATGCCTCTGCCGACAGTGGTCC |
| *GDI2-1* | |
| GDI2-Fw | CATGGGCATATTTGAGAAACGTCG |
| GDI2-Rv | CGGTGCATTCCCCTCTTGCAGAC |
| LB-TDNA-SALK | GAACAACACTCAACCCTATCTCGGGC |
| *GDI3-1* | |
| GDI3-Fw-7ex | GACTTACATGCTCAACAAGCCCGAA |
| GDI3-Fw-7in | CTCATGAACCTTCAATATCATCCCT |
| GDI3-Rv | CGCAAGTTATCAATTCTCTGCAG |
| LB-FLAGdb/FST-Taq5 | CTACAAATTGCCTTTTCTTATCGAC |
| RB-FLAGdb/FST-Taq3 | CTGATACCAGACGTTGCCCGCATAA |
